# Phosphatidic Acid Dependent Recruitment of Microtubule Motors to Spherical Supported Lipid Bilayers for *In-vitro* Motility Assays

**DOI:** 10.1101/2023.10.31.564930

**Authors:** Pankaj Kumar, Paulomi Sanghavi, Dwiteeya Chaudhury, Roop Mallik

## Abstract

A variety of membrane-bound vesicles are transported by the kinesin and dynein motor proteins respectively towards plus and minus ends of microtubules inside cells. How specific lipids, particularly rare lipids, bind and activate motors on the vesicle membrane and the biological consequences thereof are poorly understood. Here we prepare <u>*spherical*</u> supported lipid bilayers (SLBs) consisting of a latex bead enclosed within a membrane that contains phosphatidic acid (PA). When incubated with motor-protein enriched fractions from *Dictyostelium* or from Rat brain, PA-coated SLBs exhibit robust motion along *in-vitro* polymerized microtubules. We quantify the activity of SLB-bound motors by measuring their force generation against an Optical trap on polarity-labelled microtubules. Such experiments reveal that PA recruits dynein-dynactin from *Dictyostelium*, but kinesin-1 from rat brain fractions to drive SLB motion. Kinesin-1 is able to dominate over dynein when dynactin, an activator of dynein, is deficient on SLBs. We demonstrate how inhibiting only kinesin can block both kinesin and dynein activity on a cargo, thus providing an explanation for the long-standing “*paradox of co-dependence*” in the field of intracellular transport. We also find that as the size of a motor-driven SLB increases, the SLB is able to reverse direction more easily along a microtubule. This observation provides clues as to how micron-sized cargoes are efficiently able to navigate the crowded cytoskeletal space inside cells.

## INTRODUCTION

Intracellular transport of vesicular cargoes (e.g. endosomes, mitochondria, phagosomes) requires force from motor proteins that attach to the membrane of cargoes directly, or via adaptor proteins (1, 2). Biophysical function of motors has been investigated *in-vitro* using microtubule-gliding and bead-motility assays (3). In both assays, however, the motor is usually attached *sans* membrane to a non-physiological surface such as a glass coverslip or a plastic bead. How membrane lipids affect motor recruitment and activity (i.e. force), and the implications thereof, are therefore poorly understood (2). The lipid membrane is not just a passive surface, rather, it can recruit specific motors and modulate how the motor functions on the cargo. For example, the kinesin-3 family member KIF16B binds to phosphatidylinositol-3-phosphate (PI[3]P) via a PX motif whereas the KIF1A/Unc104 motor binds to phosphatidylinositol-4,5-bisphosphate (PI[4,5]P_2_) via its PH domain (4). Microtubule gliding assays suggest that cholesterol prevents slippage of kinesin against opposing force (5). A collection of myosin-V motors is able to transport synthetic liposomes faster when the underlying lipid is in a fluid state, perhaps because trailing myosins detach more easily in this situation (6). Similar mechanisms are also suggested for the kinesin-1 motor (7). Transport inside cells can also be affected profoundly by the lipid composition of an organelle. Such effects are known for the endo-phagosomal pathway (8, 9) and for lipid droplets (10). The existing literature on lipid-motor interactions and its potential implications to cell/physiological functions has been reviewed by us (2).

In the context of this study, phosphatidic acid (PA) is a rare but highly bioactive phospholipid with established functions in cell signaling (11), vesicular trafficking (10, 12–14) and physiology/lipid homeostasis (10, 15). PA is an acidic phospholipid with a small negatively charged head group and a conical shape. Thus, PA interacts with a variety of proteins (including motors) through a unique combination of electrostatic, hydrogen bonding and curvature dependent mechanisms (11, 13). In metastatic tumour cells, vesicles containing metalloproteases have PA on their membrane. This PA binds directly to the conventional kinesin (kinesin-1) motor, causing transport of these vesicles to the cell periphery, following which metalloproteases are secreted out to invade neighbouring tissue (14). We have shown that PA-dependent recruitment of kinesin-1 to lipid droplets in the liver is important for lipoprotein secretion and systemic lipid homeostasis (10, 16). Insulin signaling generates PA on the lipid droplet membrane in hepatocytes inside the liver of the animal (rat) in its fed state. Kinesin-1 binds directly to PA, transporting lipid droplets to the periphery of hepatocytes, where droplets are likely catabolized by enzymes present in the smooth endoplasmic reticulum to obtain triglycerides for lipoprotein production. This pathway is toned down when insulin signaling is diminished in the fasted state, allowing the liver to efficiently control circulating serum triglycerides in the animal (10). Taken together, PA connects vesicle transport to tumour proliferation, systemic lipid homeostasis and possibly other unknown biological pathways. We emphasize that PA is a rare phospholipid that exists at a few mole-percent in membrane lipids, but has high affinity to specific proteins (11, 13). Therefore, even small changes in PA on membranes can have profound effect on cell/physiological functions (10, 14). The above findings reveal novel disease-relevant roles for PA-motor interactions, but these are not easy to dissect using a completely *in-vivo* approach. Such studies could be advanced if certain aspects of PA-motor interactions can be reconstituted outside cells.

With this motivation, here we develop an assay to dissect PA-motor interactions *in-vitro*. We prepare artificial cargoes that consist of latex beads coated with a membrane containing phosphatidylcholine (PC) and PA. These structures are essentially “*<u>Spherical</u>* Supported Lipid Bilayers”, but will just be referred to as Supported Lipid Bilayers (SLBs). Others have studied kinesin function on spherical (7) and planar SLBs (5). However, those studies used bacterially expressed truncated kinesin motors that were attached to functionalized lipids via artificial linkages. Therefore, those experiments excluded motor-associated regulatory proteins that may affect motor function (1, 17). Further, the artificial motor-lipid linkages may modify force generation by motors/motor-complexes. To study how motors function on lipids in a more native-like context, we recruit motors to PA-containing SLBs from an “ATP releasate” that is enriched in motors and motor-associated proteins. We find that PA causes motor-driven SLB motion of distinct varieties, depending on the source of the ATP releasate. PA-containing SLBs (referred to as PA-SLBs henceforth) recruit dynein and dynactin from *Dictyostelium* releasate to drive minus-directed SLB motion, but these SLBs recruit kinesin-1 from rat brain releasate to drive plus-directed SLB motion. Irrespective of the source and type of motors, PA significantly enhances motion and force generation by motors. These inferences are supported by the use of specific inhibitors against dynein and kinesin-1 that induce surprisingly contrasting effects on SLB motion. Thus, diverse kinds of lipid-dependent SLB motion can be generated and investigated in mechanistic detail using the SLB assay developed here.

## RESULTS

### Preparation of Motor-bound SLBs for Optical Trapping and Biochemical Assays

We first prepared liposomes of a defined lipid composition doped with trace amounts of the fluorescent lipid Rhodamine-phosphatidylethanolamine (0.01% Rh-PE). The liposomes were then deposited onto a latex bead (500nm diameter) as described in Supplementary Fig 1A (also see Methods). Imaging of Rh-PE showed the SLB membrane as a ring-like structure enclosing the bead tightly in confocal sections (Supplementary Fig 1B). SLBs of different lipid composition (see below) were next incubated with ATP-releasate that was prepared from *Dictyostelium* or from Rat brain. The preparation of ATP-releasate is described (Methods). Briefly, we incubated clarified cytosol with microtubules (MTs) in the absence of ATP to allow binding of motors and MT-associated proteins to the MT. MTs were pelleted, after which ATP was added to release motors into the supernatant, thus yielding the ATP-releasate. SLBs were then incubated with ATP-releasate on ice for 5 minutes to allow attachment of proteins (including motors) to lipids on the SLB membrane. This method avoids multiple rounds of centrifugation that is required for purification of endogenous motors from mammalian cells/tissues [e.g. see Ref. (18)], during which enzymatic activity of motors can degrade.

As demonstrated extensively by us and others earlier using latex-bead phagosomes (19–21), the buoyancy of latex beads allows rapid separation of a large and highly pure fraction of SLBs that can be subjected to bulk biochemical assays to identify SLB-associated proteins. Further, the spherical refractile beads enclosed within the negligibly thin (∼5nm) membrane makes the SLB an ideal cargo for optical trap based force measurements (7, 20). SLB motion was assayed by flowing in the SLBs along with the ATP-releasate on to *in-vitro* polymerized MTs that had been adhered previously to a coverslip inside a flowcell. Individual SLBs were caught in the trap and held close to a MT. The underlying MT and SLB were visualized in a differential interference contrast (DIC) microscope equipped with an optical trap as described earlier (22). We observed force generation by SLB-bound motors against the trap, as well as free motion of SLBs after switching off the trap. The use of polarity-labelled MTs (23) allowed us to determine the activity of kinesin versus dynein in real time. These MTs have a biotinylated minus-end marked by streptavidin coated beads (Methods).

### PA Enhances Dynein driven SLB Motion with ATP Releasate from *Dictyostelium*

To probe the effect of PA, SLBs were prepared using only PC (called PC-SLBs), or with 95 mole% of PC + 5 mole % of PA (PA-SLBs). PA levels in cell membranes are between 2 to 4.4 mole% (24), and previous studies have used PC:PA in similar ratio (14, 25). SLBs incubated with *Dictyostelium* ATP releasate were held above a MT with the trap to engineer motility events, and the ensuing SLB motion was assayed. Fig 1A shows representative distance versus time tracks of the free motion of PC and PA-SLBs along MTs. The optical trap was switched off right after an SLB started to move in this experiment. Free motion of representative PC-SLBs and PA-SLBs is also shown in Supplementary Movies 1 and 2, respectively. The video tracks were parsed into segments of constant velocity (26, 27), as shown for a representative track in Supplementary Fig 2A. Based on such parsing, PA-SLBs moved with significantly faster velocities (Fig 1B). Overall, PA-SLBs exhibited qualitatively improved motion that was longer, smoother and faster.

We next assayed force generation by motors on individual SLBs against the trap on polarity labelled MTs, so that activity of plus or minus directed motors could be identified and separated. These events were recorded using a Quadrant photo detector (22) at high spatial and temporal resolution (∼5 nanometres; 1 KHz sampling frequency), making it unlikely that any transient activity of motors would be missed. Studying SLBs in an optical trap on polarity labelled MTs allows rapid, real-time measurement of the force (i.e. function) of lipid-bound kinesin or dynein motors. SLBs were separated into four categories based on their behaviour in the trap over a fixed observation window of 45 seconds per SLB, as depicted in Table-1:-

**TABLE 1.**

| TABLE 1 | SLB CATEGORY | SLB MOVEMENT AGAINST OPTICAL TRAP<br>(45 sec Observation window) |
| --- | --- | --- |
|  | *Non-motile* | Never moved. |
|  | *Bidirectional* | Towards <u>Both</u> (Plus and Minus) ends of the MT |
|  | *Plus-directed* | <u>Only</u> towards Plus-end of the MT |
|  | *Minus-directed* | <u>Only</u> towards Minus-end of the MT |

Fig 1C shows 20-seconds of optical trap data for a “plus-directed” PC-SLB and a “minus-directed” PA-SLB, so called because the given SLB showed repeated stalls against the trap in only one direction (see schematic of the MT in Fig 1C). Examples of “Bidirectional” motion are shown in Supplementary Fig 2B. Most PA-SLBs exhibited repeated minus-directed stalls in rapid succession (Fig 1C, lower panel). As also seen in Fig 1C, PA-SLBs moved farther away from the trap centre as compared to PC-SLBs, suggesting higher force generated by motors on the PA-SLB. Fig 1D shows the distribution of SLBs within the categories of Table-1, as determined from optical trapping data. Note that 34% of PC-SLBs were non-motile, but this fraction was only 9% for PA-SLBs. Thus, a high overall fraction (∼91%) of SLBs showed robust activity of motors in presence of PA. Taken together, PA significantly improved the activity of *Dictyostelium* motors on SLBs as measured by different parameters.

Unc104, a processive kinesin-3 family motor is the major plus-directed motor for long-range transport in *Dictyostelium* (4). The majority of plus-directed motion for PC-SLBs is likely driven by Unc104 (see next section). PA-SLBs, which exhibit improved overall motility, also showed a remarkable reversal in the direction of motion compared to PC-SLBs. The majority of PA-SLBs (64%) were minus-directed, with only 15% being plus-directed and 12% bidirectional (Fig 1D). Thus, a minus-directed motor is highly activated by addition of PA to SLBs. This motor is likely dynein because processive minus-directed kinesins are not found in the genome of *Dictyostelium* (28). In an earlier report, we have found that Dynein motors cluster into cholesterol-rich microdomains on mature phagosomes, yielding robust motion and frequent minus-directed stalls (9). The PA-SLB motion seen here therefore has some qualitative similarities to phagosome motion, although PA-SLBs do not contain cholesterol.

### PA recruits the *Dictyostelium* Dynein-Dynactin complex to SLBs

*Dictyostelium discoideum* ATP releasate contained Unc104, dynein and dynactin (Supplementary Fig 3A). To evaluate the affinity of these motors to lipids on the SLB, we conducted a “SLB pulldown” experiment. PC or PA-SLBs were incubated with ATP-releasate from *Dictyostelium*. The SLBs were then separated by centrifugation, washed, and probed by western blotting for lipid-bound motors. PA-SLBs recruited significantly more dynein compared to PC-SLBs, but Unc104 kinesin was recruited equally to PC and PA-SLBs (Fig 1E,1F). The SLB pulldown also detected dynactin on PA-SLBs, but not on PC-SLBs (Fig 1E).

**Figure 1.**
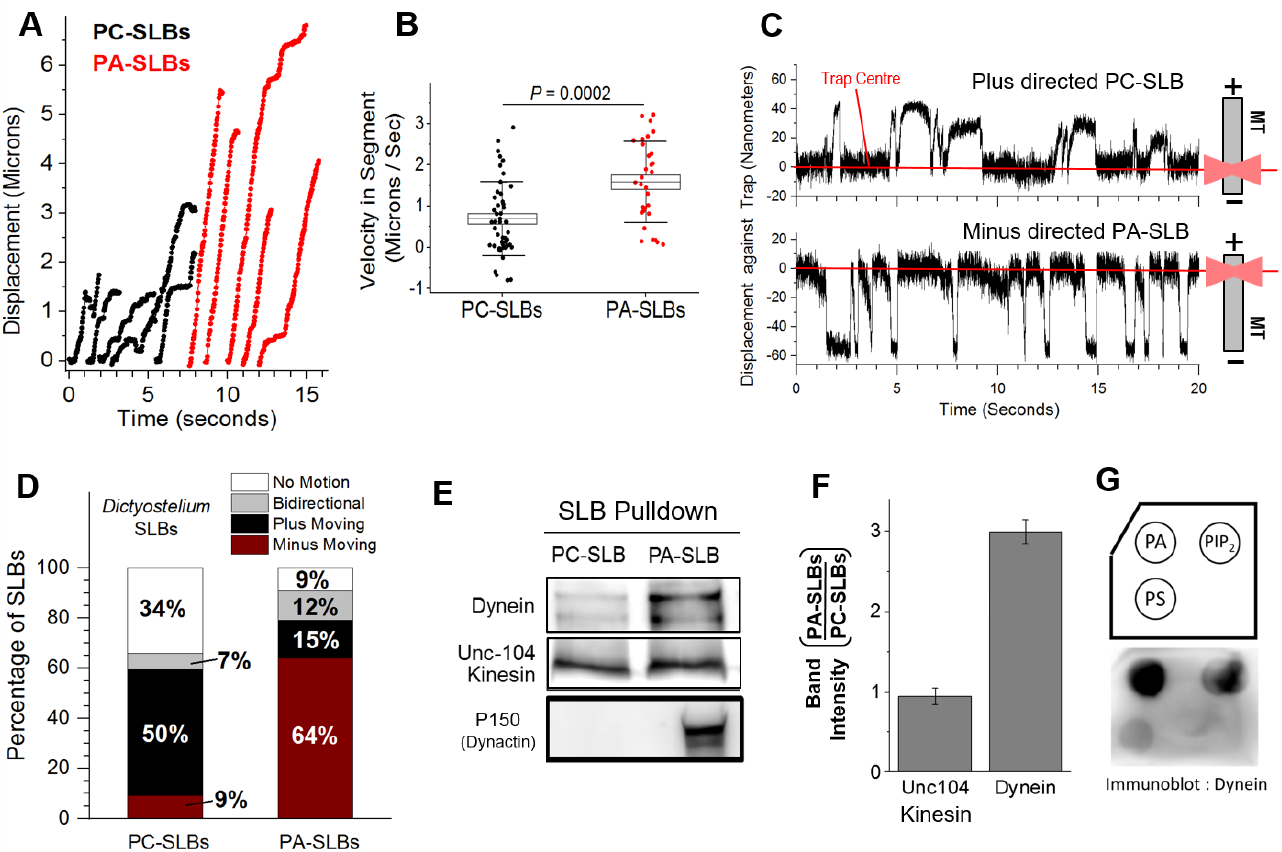
Phosphatidic Acid Enhances Minus-directed Motion of SLBs Prepared using *Dictyostelium* ATP releasate. A. Representative tracks for free runs of PC-SLBs (black) and PA-SLBs (red) shown as a displacement versus time graph. The direction of motion (towards plus or minus end of MT) is not specified for ease of representation. B. Box plot representing the velocity within constant-velocity segments for PC and PA-SLBs (see main text). PA-SLBs have higher average velocity than PC-SLBs. C. Displacement vs. time graph for a PC-SLB moving toward plus and PA-SLB moving toward minus direction against an optical trap. Schematics on the right depict the experimental geometry and MT polarity. Red lines indicates centre of the trap (Zero position). These direction-specific tracks are representative of the dominant direction of motion for PC-SLBs (plus) and PA-SLBs (minus). The PA-SLB shows more frequent force generation events as compared to the PC-SLB. The PA-SLB also moves out farther (∼60nm), and therefore motors on the PA-SLB generate higher force compared to PC-SLB (∼40nm). D. Categorization of SLBs based on their motion on polarity-labeled MTs. Very few PA-SLBs are immotile as compared to PC-SLBs. Note the drastic increase in minus-moving SLBs upon addition of PA. A total of 32 PC and 95 PA-SLBs were analysed. E. Western blots representing motors retained on SLBs after an “SLB pull down”. Number of PA-SLBs and PC-SLBs were normalized using the Optical density at 500nm (methods). PC and PA-SLBs were incubated with *Dictyostelium* ATP releasate, washed, and probed for *Dictyostelium* Dynein, *Dictyostelium* Unc104 kinesin and *Dictyostelium* Dynactin. F. Quantitation of motor binding to PC and PA-SLB. PA has no significant effect on Unc104 recruitment to SLBs. However, Dynein amount on PA-SLBs is ∼3 times of that bound to PC-SLBs. (Mean ± SEM, N=3) G. Lipid Blot to show that negatively charged lipids such as Phosphatidic Acid (PA) and Phosphatidylinositol-4,5-bisphosphate (PIP2) can recruit *Dictyostelium* Dynein. Phosphatidylserine (PS) shows poor recruitment of Dynein.

The interaction between dynein and dynactin, and therefore the motion driven by dynein-dynactin complexes, can be disrupted by a peptide that mimics the dynein intermediate chain (29, 30). We have cloned the dynein intermediate chain from *Dictyostelium* and shown that this peptide specifically inhibits minus-directed motion of SLBs prepared from *Dictyostelium* ATP releasate (20). Taken together, these findings support the conjecture that minus directed PA-SLB motion is driven by the *Dictyostelium* dynein-dynactin complex. However, while 50% PC-SLBs were plus-directed, very few (only 15%) PA-SLBs were plus-directed (Fig 1D). It therefore appears that the excess dynein-dynactin recruited to PA-SLBs confers minus-directed motility to PA-SLBs by dominating over Unc104. This may be expected because the Unc104 kinesin driven motion of PC-SLBs appears irregular compared to PA-SLBs (Figs 1A, 1C).

To better understand lipid-motors affinities, we next performed lipid blot assays using *Dictyostelium* ATP releasate. PA is an acidic phospholipid with a negatively charged head group and a conical shape (11, 13). While the other acidic phospholipids such as phosphatidylserine (PS) and phosphatidylinositol-4,5-bisphosphate (PIP_2_) also bound to *Dictyostelium* dynein, their affinity was significantly lower than PA (Fig 1G). We therefore believe that in addition to its negative charge, the conical shape of PA also has a role in dynein-dynactin binding. Taken together, this data suggests that PA-SLBs can be used to recruit dynein-dynactin from *Dictyostelium* for analyzing dynein function on a lipid membrane. PC-SLBs, which are largely driven towards the plus end of the microtubule, may also be used to investigate the Unc104 kinesin motor. Note that Unc104 kinesin recruitment is insensitive to PA, whereas dynein-dynactin recruitment depends on PA (Fig 1E). Therefore, it should be possible in future to tune the ratio of Dynein:Kinesin on SLBs simply by varying the amount of PA on SLBs. Such experiments can help to understand the competition between opposing motors as they drive a cargo (31).

### The Source Matters : PA Enhances Kinesin-1 driven SLB Motion with Rat brain ATP*-*Releasate

Keeping in mind the well-established functions of PA in cell signaling (11), vesicular trafficking (10, 12–14) and physiology/lipid homeostasis (10, 15), we next probed PA-motor interactions in an evolutionarily distant system, the mammalian brain. ATP-releasate was prepared from rat brain and incubated with PC or PA-SLBs (Methods). Fig 2A shows representative video tracks of the free motion (optical trap switched off) of PC and PA-SLBs. The motion of PC and PA-SLBs can also be seen in Supplementary Movies 3 and 4 respectively. Video tracks were parsed as described earlier to obtain the distribution of segment velocities (Fig 2B). Unlike *Dictyostelium-*SLBs, no significant increase in velocity was seen for PA-SLBs incubated with rat brain releasate (compare Figs 2A,2B with Figs 1A,1B).

We next moved to optical trapping of SLBs on polarity labelled MTs. Fig 2C shows representative data for a minus-directed PC-SLB, a plus-directed PA-SLB and also a minus-directed PA-SLB. As described in Table 1, we separate the observed motion into four categories and depict these categories in a bar graph (Fig 2D). Note that 42% of PC-SLBs are non-motile, but this number is only 10% for PA-SLBs. PA causes this improvement in overall motility by increasing the plus-directed fraction of SLBs (8% for PC, but 42% for PA-SLBs). However, PA has no effect on minus-directed SLB motion (42% for PC, 43% for PA-SLBs), a finding that contrasts with the PA-induced improvement of minus-directed motion for *Dictyostelium-*SLBs (Fig 1D). PA therefore improves overall SLB motility for both rat brain releasate and *Dictyostelium*, but this improvement has different mechanisms. For rat brain, plus-directed (kinesin driven) motion is upregulated, whereas for *Dictyostelium* minus-directed (dynein driven) motion is upregulated. An obvious explanation for this difference could be that more kinesin is recruited from rat brain releasate to PA-SLBs. To investigate the PA-to-Kinesin interaction on SLBs, we did an SLB pull-down with rat brain ATP releasate. A small, but consistent increase of kinesin-1 was observed on PA-SLBs (Figs 2E, 2F), which may be expected as kinesin-1 binds directly to PA (10, 14). Dynein was recruited equally to PC and PA-SLBs (Fig 2E and 2F), consistent with the equal fraction of minus-moving PC and PA-SLBs (Fig 2D). A lipid blot supported the possibility that kinesin-1 in rat brain releasate has higher affinity for PA (Fig 2G). We have earlier found that PA has no detectable affinity for kinesin-2 or dynein from mammalian cell lysate (25).

Further, we noted that Dynactin was not detectable in rat brain ATP releasate (Supplementary Fig 3B), an observation that agrees with others (32). Accordingly, dynactin was also barely detectable on PA-SLBs prepared from rat brain releasate (Fig 2E). Dynactin is therefore present abundantly on PA-SLBs prepared from *Dictyostelium* (Fig 1E), and is functional on SLBs as shown by specific inhibitors (20), but dynactin is likely deficient on the PA-SLBs prepared from rat brain ATP releasate. Taken together, the enhanced plus-directed motion for PA-SLBs with rat brain releasate likely derives from two factors. First, somewhat higher abundance of kinesin-1 on PA-SLBs (Figs 2E,2F). Second, deficiency of dynactin (which is an activator of dynein) in rat brain releasate, and therefore also on PA-SLBs.

**Figure 2.**
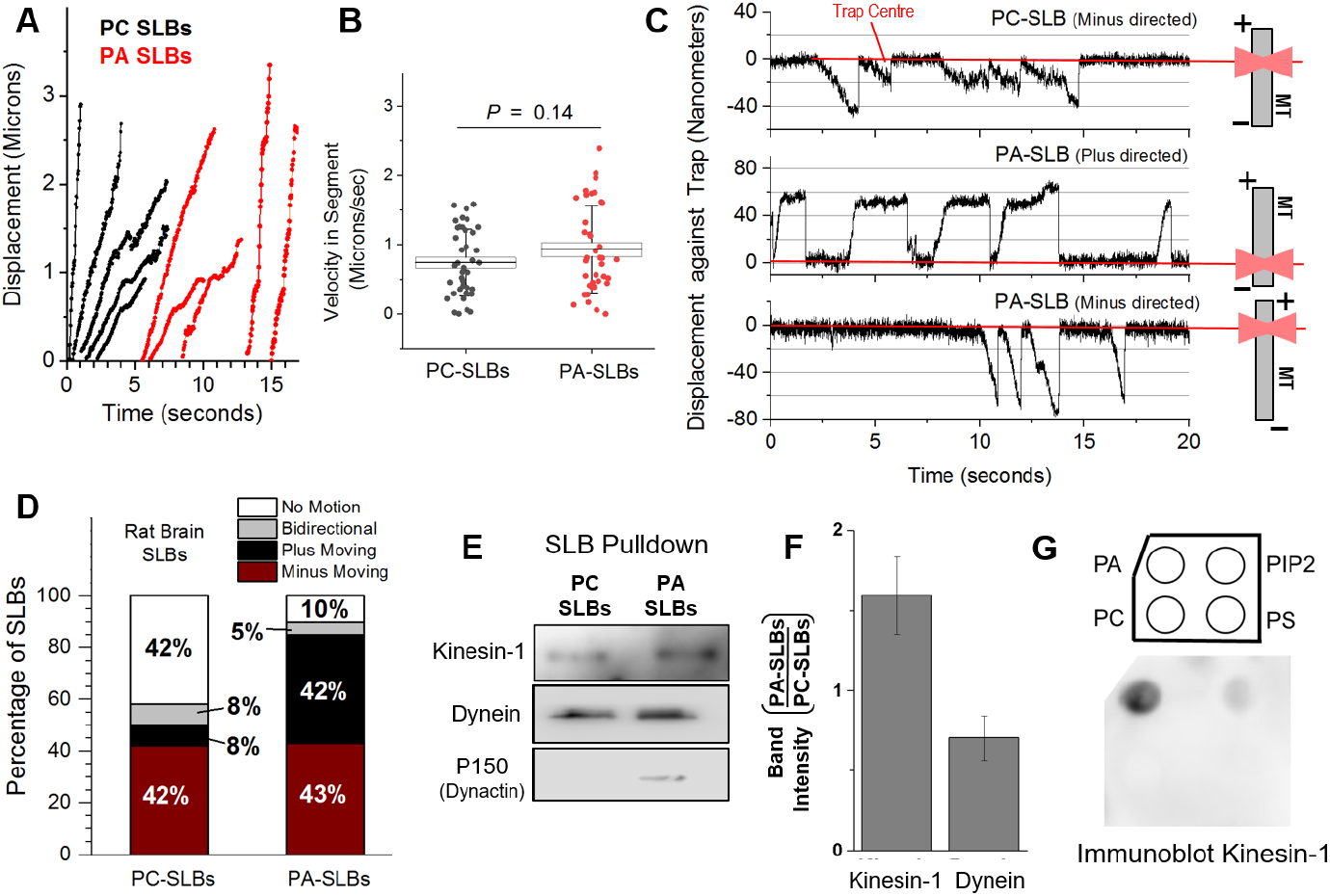
Phosphatidic Acid enhances Plus-directed Motion of SLBs prepared with Rat brain ATP releasate. A. Representative tracks for free runs of PC-SLBs (black) and PA-SLBs (red) shown as a displacement versus time graph. The direction of motion (towards plus or minus end of MT) is not specified for ease of representation. B. Box plot representing the velocity within constant-velocity segments for PC and PA-SLBs (see main text). There is no significant difference in the velocities. C. Displacement versus time curves for the motion of PC and PA-SLBs against an optical trap. Examples of plus as well as minus-directed motion are shown for PA-SLBs. Schematics on the right depict the experimental geometry and MT polarity. Red lines indicates centre of the trap (Zero position). D. Categorization of SLBs based on their motion on polarity-labeled MTs. PA-SLBs have a small fraction of immotile SLBs (10%) as against PC-SLBs (42%). Note the increase of plus moving SLBs upon inclusion of PA. 44 and 62 PC-SLBs and PA-SLBs were analyzed, respectively. E. Immunoblots representing motors retained on SLBs in an SLB pull-down assay carried out from the rat motor pool. SLBs were incubated with rat brain ATP releasate, washed, and probed for bound/retained motors. PC and PA-SLB numbers were normalized by optical density. F Quantitation of motors bound to PC and PA-SLBs in the SLB pull-down assay. Error bars are Mean ± SEM, N=3. A small increase in recruitment of Kinesin-1 to PA-SLBs is seen, as compared to PC-SLBs. G. Lipid Blot demonstrating the ability of negatively charged lipids to recruit Kinesin-1 from the ATP releasate fraction. Phosphatidic Acid (PA) shows strong binding of Kinesin-1. Phosphatidylinositol-4,5-bisphosphate (PIP2) shows weak recruitment of Kinesin-1. Phosphatidylserine (PS) and Phosphatidylcholine (PC) shows no detectable binding.

### Distinct Effects of Kinesin-1 and Dynein Inhibition on SLBs prepared Using Rat Brain ATP Releasate

PA-SLBs incubated with rat brain ATP releasate exhibited equal probability of plus and minus motion (Fig 2D). We therefore used these PA-SLBs to check if specific inhibitors of dynein and kinesin-1 can block one versus the other kind of motion. Inhibiting one kind of motor may or may not allow the opposing motor to take over, depending on the “Tug-of-war” Versus “Coordination” models of bidirectional transport (31, 33, 34). To specifically inhibit kinesin-1 on PA-SLBs, we used a GST-tagged kinesin tail domain peptide (KTD-GST) purified from bacteria (10). KTD-GST contains the C-terminal end of the kinesin-1 heavy chain (amino acid 854 to 963), and is known to inhibit kinesin-1 ATPase activity (35–37).

While only 10% of GST-control treated PA-SLBs were non-motile, KTD-GST used at 170 nM concentration drastically increased the non-motile fraction to 81% (Fig 3A). To effect an 81% block in motion, KTD-GST must somehow be able to inhibit motion of kinesin-driven (42% in control) as well as dynein-driven PA-SLBs (43% in control). KTD removed neither kinesin-1 nor dynein from PA-SLBs (Fig 3B, 3C). KTD is not a generic inhibitor of motors, as it had no effect on motion of PA-SLBs prepared from *Dictyostelium* releasate (Supplementary Fig 3C). Thus, KTD must be able to also somehow block dynein when it inhibits kinesin-1 on PA-SLBs. Coy *et*.*al*. (37) found that KTD reduces the velocity of MT-gliding by kinesin-1, and thus suggested that KTD reduces kinesin’s OFF-rate from the MT (35–37). Very interestingly, we observed that almost all of the non-motile SLBs in presence of KTD could not be pulled away from the MT using the optical trap. Further, KTD marginally increased the amount of kinesin-1 (but not of dynein) pelleting with MTs (Fig 3D, 3E). This assay was done in presence of ATP (∼1mM) to resemble the motility experiment, where ATP is present. Taken together, KTD appears to block kinesin as well as dynein-driven SLB motion because it causes kinesin-1 to bind more tightly (i.e. to get “locked”) to the MT.

We next used Ciliobrevin D, a well-known inhibitor of dynein (38) on PA-SLBs prepared from rat brain ATP releasate. Ciliobrevin used at 20μM expectedly reduced the fraction of minus-directed PA-SLBs (Fig 3F). However, ciliobrevin treated PA-SLBs also exhibited a surprising and dramatic compensatory increase in plus-directed motion, so that their overall motile fraction was comparable to untreated PA-SLBs (∼90%; Fig 3F). Ciliobrevin obviously did not inhibit kinesin-1 because ciliobrevin treated PA-SLBs displayed abundant plus-directed motion. Earlier reports have also demonstrated that kinesin-1 activity is unaffected upto 100μM ciliobrevin (39). Ciliobrevin did not remove dynein or kinesin-1 from PA-SLBs, as determined by an SLB pull-down experiment (Fig 3G and 3H). Unlike the effect of KTD, ciliobrevin treated PA-SLBs could be pulled off easily from MTs by the trap. Ciliobrevin also had no effect on kinesin-1 or dynein pelleting with MTs in presence of 1mM ATP (Fig 3I, 3J), suggesting that ciliobrevin is not causing dynein to bind tightly to MTs. Based on these observations, kinesin-1 appears to overcome the ciliobrevin-inactivated dyneins and cause overwhelmingly plus-directed motion of PA-SLBs. This compensatory increase of plus-directed motion by ciliobrevin contrasts with the overall arrest of PA-SLB motion after KTD treatment (compare Figs 3A and 3F). Different kinds of motor inhibition could therefore have contrasting effects on SLB motion, depending on the exact mechanism of inactivation (see discussion).

### Repeated Engagements of Same Motor-Type on SLBs : Implication to Intracellular Transport

Many kinds of vesicles move in a bidirectional (back-and-forth) manner along MTs inside cells (31, 33, 34). Early Phagosomes (EPs) prepared by phagocytosing latex beads into *Dictyostelium* cells show rapid alternating activity of dynein and kinesin in *in-vitro* optical trap experiments that were conducted similar to this study (40). Such “bidirectional” EP stalls can be seen in Supplementary Fig 2C, where once a given class of motors on the EP detaches, the opposing motors usually generates force within ∼1 second. We were therefore very surprised to find that a given SLB would almost always move in the same direction (plus or minus) in our optical trap assay. This rarity of bidirectional motion was true for all combinations of lipids and motors on SLBs (Figs 1D, 2D). The “unidirectional” nature of SLBs was most intriguing for PA-SLBs driven by rat brain ATP releasate, because as a population, these SLBs showed equal amount of plus and minus motion (Fig 2D). Both plus and minus motors therefore have almost equal activity on PA-SLBs. However, once a given motor or type of motor attaches to MT from the SLB, it continues to attach/detach/re-attach, whereas the other type of motor is not able to engage.

**Figure 3.**
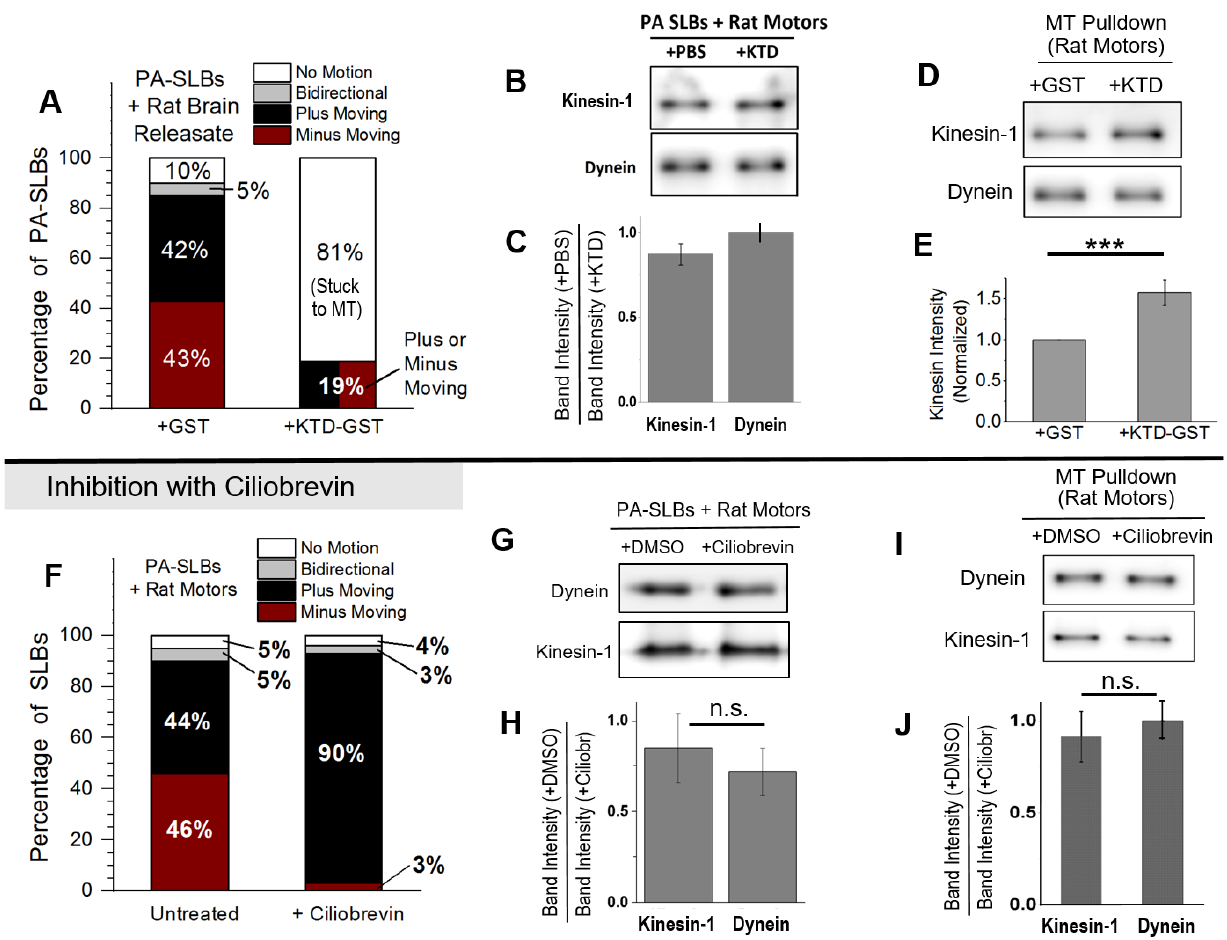
Inhibition of SLB motion prepared from Rat Brain ATP Releasate using Kinesin Tail Domain and Cilobrevin. A. Categorization of PA-SLBs incubated with rat brain releasate in presence of 170 nM GST (control) or 170 nM KTD-GST. KTD implies “Kinesin-1 tail domain”. Total of 30 and 47 SLBs were analyzed for GST and KTD-GST respectively. KTD-GST induces most PA-SLBs (72%) to get stuck on MTs. Once stuck, these SLBs cannot be removed from the MT by the trap. Increased affinity to MTs in presence of KTD-GST is also reflected in the MT pelleting assay (Fig. 3E and 3F). B. Immunoblots representing proteins retained on PA-SLBs during SLB pull-down assay. PA-SLBs were incubated with rat brain motor fractions along with PBS (as a control) or 170 nM KTD-GST, washed, and then probed for kinesin-1 and dynein. C. Quantitation of Kinesin-1 and dynein intensity in the SLB pull-down assay. No significant depletion of Kinesin-1 occurs from the SLBs upon KTD treatment. N=3. Data represents Mean ± SEM, *P* > 0.05. D. Immunoblot depicting kinesin-1 and dynein bound to microtubules after a microtubule-pelleting assay performed in presence of ATP releasate from rat brain. MT pulldown was done in presence of ATP to mimic the conditions in motility assay. GST-peptide (control) or KTD-GST was present at 17μM concentration during the MT-pulldown. E. Quantification of Kinesin-1 bound to microtubule pellet. Kinesin-1 shows significantly higher binding in presence of KTD-GST as compared to the GST-control. *N*=6. Data represents Mean ± SEM. ***p < 0.001. F. Categorization of PA-SLBs incubated with rat brain releasate, based on their motion on polarity-labeled MTs in presence and absence of Ciliobrevin. Motile fraction of PA-SLBs treated with 20μM Ciliobrevin remains unchanged (∼ 95% in each case). Ciliobrevin almost completely inhibits minus-directed motion. PA-SLB motion in presence of Ciliobrevin is almost completely plus-directed driven by Kinesin. 130 and 18 SLBs were analyzed in Ciliobrevin and DMSO groups, respectively. N=3. G. Immunoblots representing proteins retained on PA-SLBs after SLB pull-down. PA-SLBs were incubated with rat brain motor relesate in presence of 20μM Ciliobrevin or DMSO, washed, and probed for bound/retained proteins. H. Quantitation of band intensity of motors in the SLB pull-down assay. No significant depletion of motors is seen on SLBs after Ciliobrevin treatment. N=3. Data represents Mean ± SEM. I. Immunoblot depicting kinesin-1 and dynein bound to microtubules after a microtubule-pelleting assay performed in presence ATP releasate from rat brain. Experiment was done in presence of ATP to mimic the conditions in motility assay. Lanes represent data where DMSO (control) or 20μM Ciliobrevin was added. J. Quantitation of Kinesin-1 and Dynein amounts bound to microtubules in the presence of 20μM Ciliobrevin or DMSO in MT pelleting assay. Data represents Mean ± SEM, n.s. represents p > 0.05, N=3.

**Figure 4.**
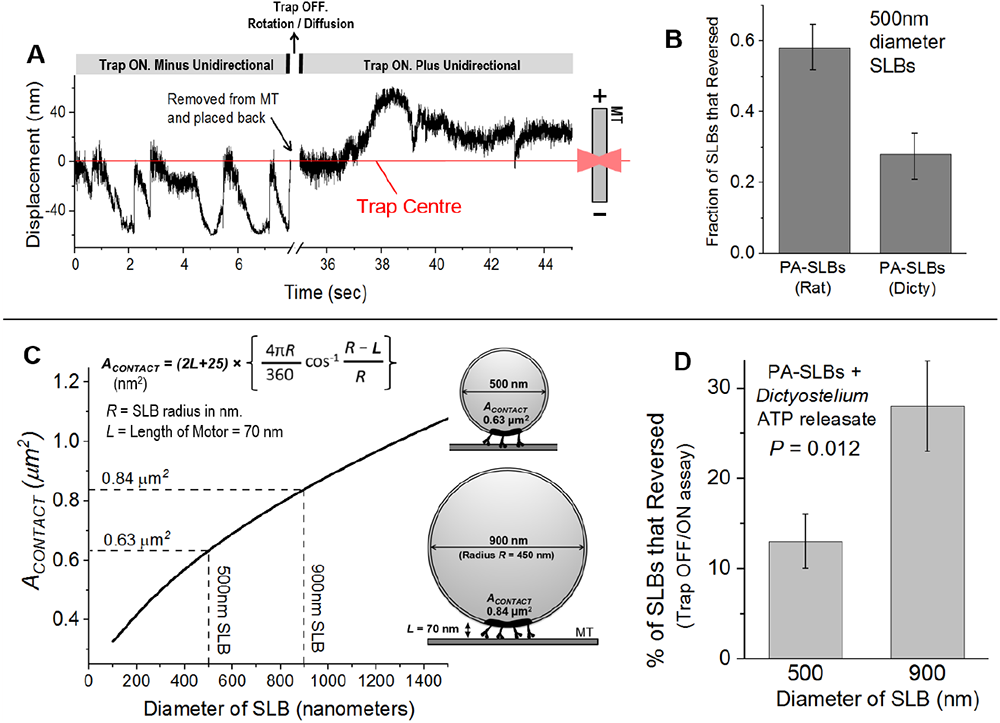
Engagement of Opposite polarity motors on SLBs. A. Displacement versus time against an optical trap for a PA-SLB prepared from rat brain releasate. The SLB initially shows dynein-driven (minus-directed) motion on a polarity labelled MT. The SLB is removed from the MT at the timepoint indicated, and then placed back on the MT after a few seconds. This separation from the MT allows rotational diffusion of the SLB. Direction of motion reverses when the SLB is placed back on the MT, presumably because a new area of the SLB (now containing kinesin-1 motors) is now proximal to the MT. B. Fraction of SLBs that reversed direction similar to the example in Fig 4A. PA-SLBs driven by rat motors show about 60% reversal probability. Probability of reversal is reduced to 25% for PA-SLBs driven by *Dictyostelium* motors. Error bars are SEM, *N* = 3. C. Cartoon insets are schematics representing *A*_*CONTACT*_, which is the region on a spherical cargo from where a motor of given size (*L* =70 nm) can access the underlying MT. The expression for *A*_*CONTACT*_ is shown. See *Sanghavi et al Current Biology 2018* for more details. The two cargoes of different size and their attached motors are drawn to relative scale. Graph represents change in *A*_*CONTACT*_ with diameter of the spherical SLB. D. Fraction of PA-SLBs driven by *Dictyostelium* motors that reverse direction on the MT when moving against an optical trap. Probability of reversal is higher for the larger cargo. Error bars are S.D. *N* = 3.

What are reversals so rare for SLBs, and might this provide interesting clues about regulation of intracellular transport? A possible, but unlikely explanation for unidirectional PA-SLBs is that some unknown mechanism causes plus and minus motors to segregate onto two separate SLB populations. A given PA-SLB with minus-directed motors may then have almost no plus-directed motors and *vice-versa*, thus yielding unidirectional motion for each population. We devised a sequence of events on a trapped SLB to test this unlikely possibility (Fig 4A). We allowed a PA-SLB to generate force, then removed the trapped SLB away from the MT and the coverslip. We now switched the trap OFF to allow the SLB to diffuse/rotate, switched the trap ON to again trap the SLB, then lowered the SLB onto the same MT for motor re-attachment. As shown in Fig 4B, ∼50% of the re-attached PA-SLBs “reversed” to now generate force only in the opposite direction (see the example in Fig 4A). The remaining 50% did not “reverse”, and continued to generate force only in the original direction. This 50-50 distribution is in good agreement with the equal fraction of plus and minus moving PA-SLBs in presence of rat brain ATP releasate (Fig 2D). Thus, a given PA-SLB incubated with rat brain ATP releasate has equal activity of both kinds of motors, *but once one type of motor engages, the other usually does not get a chance*. Does this situation hold true for PA-SLBs prepared using *Dictyostelium* ATP releasate? Indeed, such SLBs showed only ∼25% chance of reversing after they were re-attached to the MT (Fig 4B). This is exactly what one would predict because only a minor population of the *Dictyostelium* PA-SLBs show kinesin activity (15% + 12% = 27%; Fig 1D), with most of them being minus-directed (∼64%; Fig 1D).

We have earlier derived an expression for the contact area on a spherical cargo (= *A*_*CONTACT*_), from where a motor of given size can physically reach a MT (40). The expression for *A*_*CONTACT*_ is shown, and the relevant terms explained in Fig 4C, where cargoes of 500 and 900nm diameter are also drawn to relative scale. Each cargo is attached to a MT at its base via individual motors (∼70nm length). Only those motors that reside within *A*_*CONTACT*_ (black patch) are able to reach the MT. Motors outside *A*_*CONTACT*_ are not shown for the sake of clarity. Figure 4C also depicts the calculated dependence of *A*_*CONTACT*_ on cargo size. The 900nm SLB has larger *A*_*CONTACT*_ because its radius of curvature is smaller (flatter surface). Thus, the probability that plus and minus motors can both contact the MT at the same time should increase for larger SLBs, assuming that the surface density of membrane-bound motors is independent of SLB size. We therefore prepared PA-SLBs using 500nm (our original size) and 900nm diameter beads, incubated these with rat brain ATP releasate, and tested motion against the optical trap. Indeed, 900nm PA-SLBs displayed more bidirectional events as compared to 500nm PA-SLBs (Fig 4D). We estimate an increase of ∼33% in *A*_*CONTACT*_ [= 100 x (0.84 − 0.63)/0.63] going from a 500 to a 900nm SLB, but the increase in bidirectional motion is much larger at ∼115% [= 100 x (28 − 13)/13]. It therefore appears that larger cargoes are significantly more prone to reversals during motion.

## DISCUSSION

The assay using spherical SLBs is unique because it allows Optical trap based measurement of motor protein activity at very small PA concentrations, which is in the physiologically relevant range for PA. As an example, we have measured activity of motors on membranes containing only 5 mole% PA. Such biophysical measurements using SLBs may therefore help dissect how rare lipids activate motors to generate polarized distributions of vesicles inside cells for downstream cellular requirements. We found that PA recruits high levels of dynein-dynactin on SLBs from *Dictyostelium* releasate. Such PA-SLBs show robust, long-distance minus directed motion that is qualitatively similar to late phagosomes (9). Dynein motors are clustered into cholesterol enriched microdomains on the phagosome, but such clustering is unlikely to be induced by PA because it is present at very small concentrations. It is possible that being a conical lipid, PA generates negative curvature to locally raise the SLB membrane (13), and this allows PA-bound dyneins to access the MT more easily. In contrast to *Dictyostelium*, PA-SLBs exhibit kinesin-1 driven plus-directed motion when incubated with rat brain ATP releasate. This may derive from the deficiency of dynactin on such PA-SLBs, because of which kinesin-1 is able to overpower dyneins once it gets recruited via PA. Thus, dynactin could have an important role in tilting the balance in favour of dynein during a competition with kinesin. This conjecture is further supported by our *Dictyostelium* data where the highly processive Unc104 kinesin is equally present on PC and PA-SLBs (Fig 1E), yet the appearance of dynein-dynactin on PA-SLBs significantly suppresses plus-directed motion (Fig 1D). This requirement of dynactin for dynein to compete against kinesins is particularly interesting because neurodegenerative mutations in dynactin reduce the processivity of dynein-dynactin (32), and therefore possibly its ability to compete against kinesin.

Because PA-SLBs prepared with rat brain releasate exhibit equal probability of plus and minus motion, we used this system to understand how inhibitors of specific motors affect cargo motion. Very interestingly, KTD (a kinesin-1 inhibitor) reduced both kinesin-1 and dynein driven motion, whereas ciliobrevin (a dynein inhibitor) blocked only dynein with a compensatory increase in plus-directed motion (Fig 3A, 3F). Almost all KTD treated non-motile SLBs could not be pulled away from the MT by the trap. We therefore believe that KTD blocked the ATPase activity of kinesin-1, causing SLBs to get “locked” onto MTs and thus also blocking dynein activity. However, ciliobrevin inactivated dynein without a locking effect, and kinesin-1 could now dominate the SLB motion (Fig 3F). Thus, KTD and ciliobrevin have contrasting effects on SLB motion. This finding has specific implications to the *“Paradox of co-dependence”*, an unexplained phenomenon wherein inhibiting one motor-type (dynein or kinesin) enhances motion in the opposite direction in some instances, but in other instances can also block both motors to completely abrogate motion. This paradox has been discussed in detail, and numerous inhibition based studies have been cited in Ref. (31). Based on our results with SLBs, the *paradox of co-dependence* can be explained if motor-inhibitors can be classified as “Locking Inhibitors” (e.g. KTD) or “Non-locking Inhibitors” (e.g. Ciliobrevin).

PA-SLBs incubated with rat motors showed motion in both directions, but a given SLB rarely reversed direction. However, if this SLB was pulled away from the MT and then replaced back, then it could move in the opposite direction (Fig 4A, 4B). Therefore, plus as well as minus directed motors are both present on a given PA-SLB. However, once it starts moving, a given SLB only generates force in direction. It appears that such a moving SLB cannot rotate easily because it is bound to the MT by motor(s), and also because the surface (e.g. coverslip) increases the rotational drag on the SLB. It also appears that diffusion of motors within the SLB membrane is not sufficient to bring opposite-polarity motors within *A*_*CONTACT*_. Increasing SLB size promoted reversals (Fig 4D), likely because this increased *A*_*CONTACT*_ to allow both kinds of motors to bind MTs from the SLB at the same time or in quick succession. Thus, a larger cargo may be able to back out and resume motion in a different direction when facing obstructions in the crowded cytoskeletal space. If this is indeed true, it raises the question of how small endosomes/phagosomes are able to exhibit highly bidirectional (saltatory) motion (34). This question is significant because such saltatory motion is likely required for recycling of endocytosed material (e.g. transferrin) to the plasma membrane (41). Perhaps the complex lipid composition of the endosomal membrane ensures a homogeneous distribution of diverse lipid species, and therefore of kinesins and dyneins bound to these lipids. Once this happens, both motors may reside within *A*_*CONTACT*_ even on small endosomes. Such possibilities could not be replicated in our two-component (PC and PA) SLB membrane. However, the SLB assay holds promise to explore these effects because one can use a native-like lipid mixture such as the Folch fraction from brain to make SLBs, and then study their motion in presence of ATP releasate. Super-resolution imaging may permit visualization of lipid-dependent motor distribution on complex membranes to explain how lipid-motor interactions promote one or more kinds of motors to be functional for vesicle transport.

## Acknowledgements

We thank the Tata Institute of Fundamental Research Mumbai (TIFR) for providing certain resources for our experiments. Particularly, we thank Dr. Vidita Vaidya and Dr. Shital Suryavanshi for acces to the animal facility at TIFR.

