## Supplementary Information for "Phosphatidic Acid Dependent Recruitment of Microtubule Motors to Spherical Supported Lipid Bilayers for *In-vitro* Motility Assays"

*Pankaj Kumar et. al.*

#### SUPPLEMENTARY INFORMATION

#### Includes

**Supplementary Figures 1,2,3**

**Methods**

**Additional References in Methods**

**Legends for Supplementary Movies 1,2,3,4**

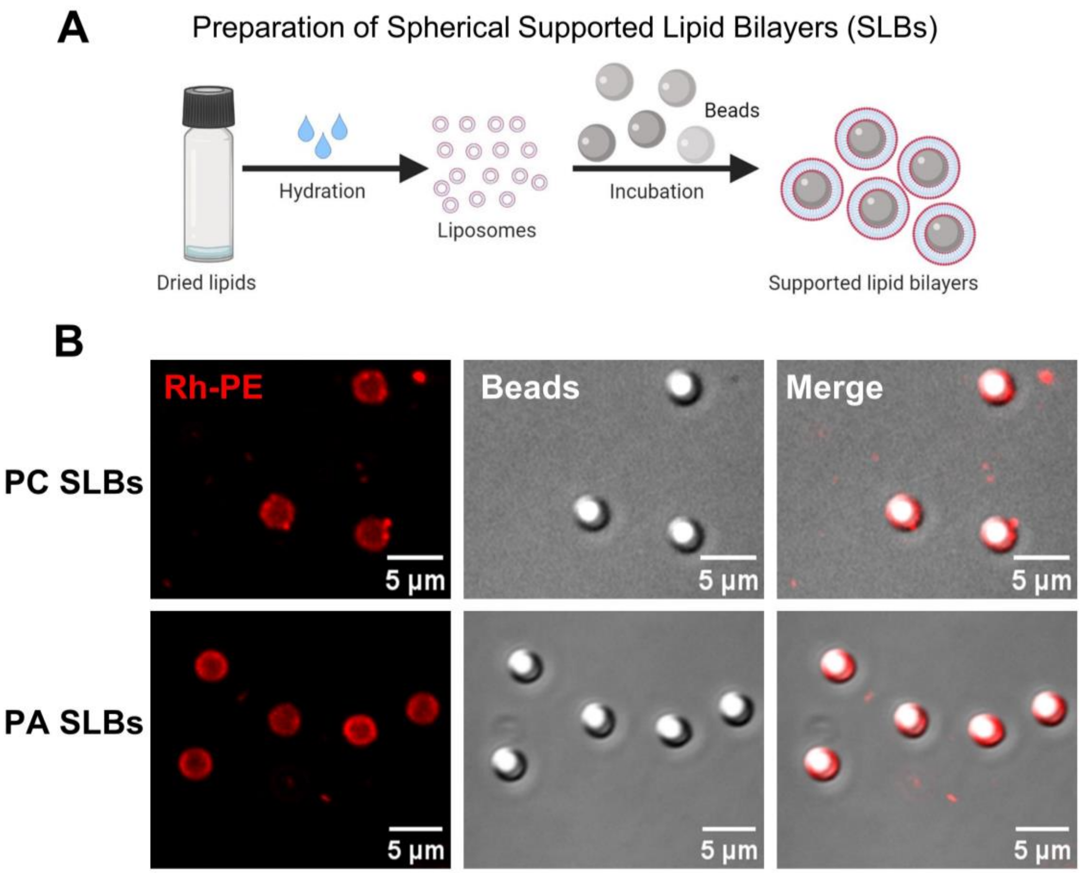


#### Supplementary Figure 1

A. Schematic for preparation of liposomes and Spherical Supported Lipid Bilayers (SLBs) from liposomes. Briefly, lipid mix of the desired composition is dried into a lipid film, hydrated, and then incubated with beads to form SLBs. The buoyant SLBs are then separated by centrifugation.

B. Confocal image of 3µm (diameter) PC-SLBs and PA-SLBs. The red signal (left-most panel and merge) is Rhodamoine-PE, present in 0.01-mole percentage to visualize the lipid membrane.

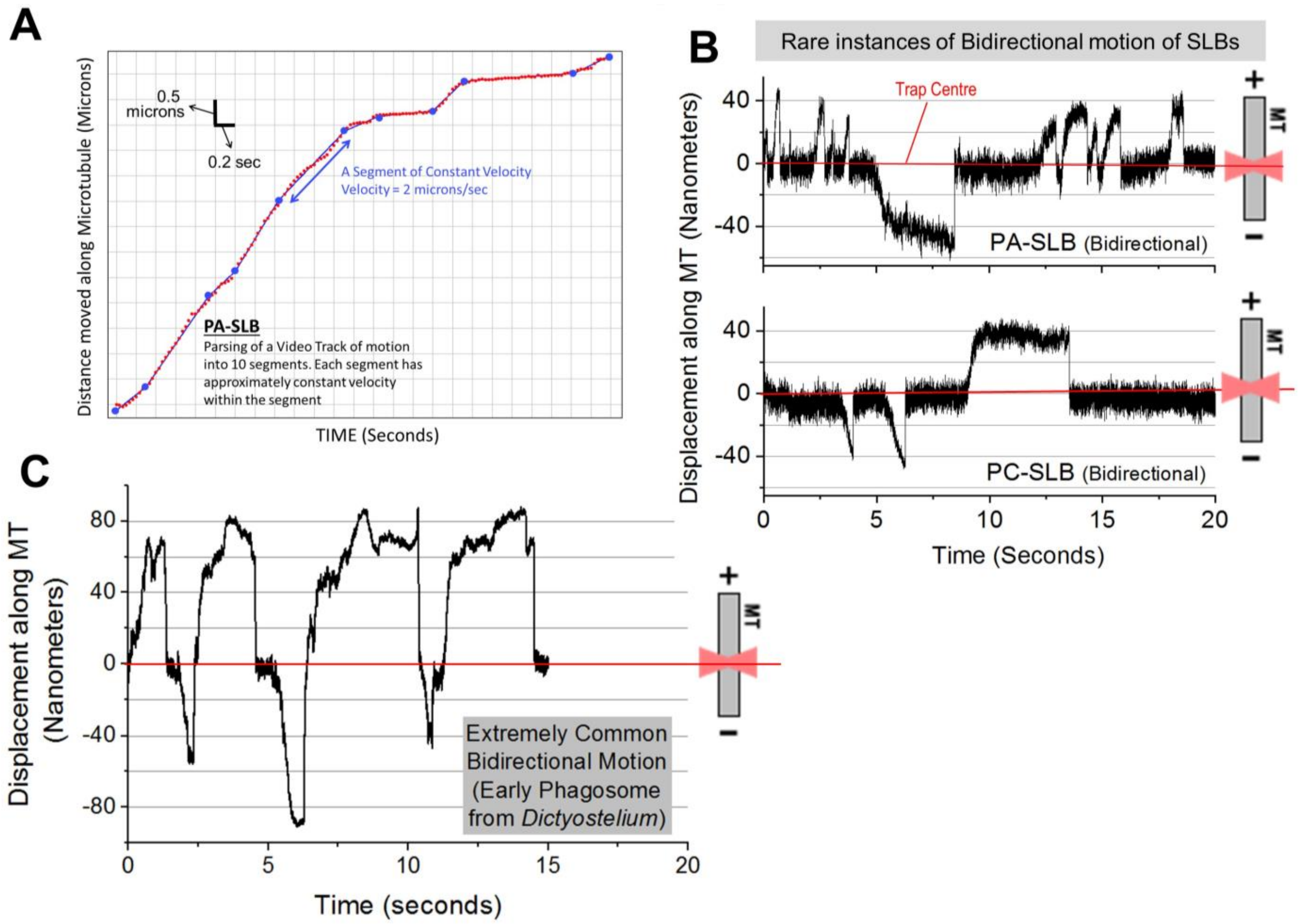


#### Supplementary Figure 2

A. Distance versus time track (red dots) of a PA-SLB moving freely along a MT (trap switched OFF). A Bayesian algorithm is applied to parse this track into 10 segments, each having an approximately constant velocity within the segment. The parsed segments are shown as large blue dots connected by a blue line. One such segment having a velocity (= slope) of 2 microns/sec is shown. The velocities obtained from such parsing are used in Figs 1B and 2B.

B. Bidirectional displacements against an optical trap for a PA-SLB (top) and a PC-SLB (bottom) incubated with *Dictyostelium* ATP releasate. These are the rare incidents where a given SLB showed excursions in both directions against the trap. Red line represents the trap center.

C. Displacement-time graph representing motion against the optical trap of an “Early Phagosome” isolated from *Dictyostelium*. Unlike rare instances of bidirectional motion in SLBs (see panel B), almost all Early Phagosomes show rapid and consecutive stalls in opposite directions (For more details, see *Sanghavi et al Current Biology 2018*.

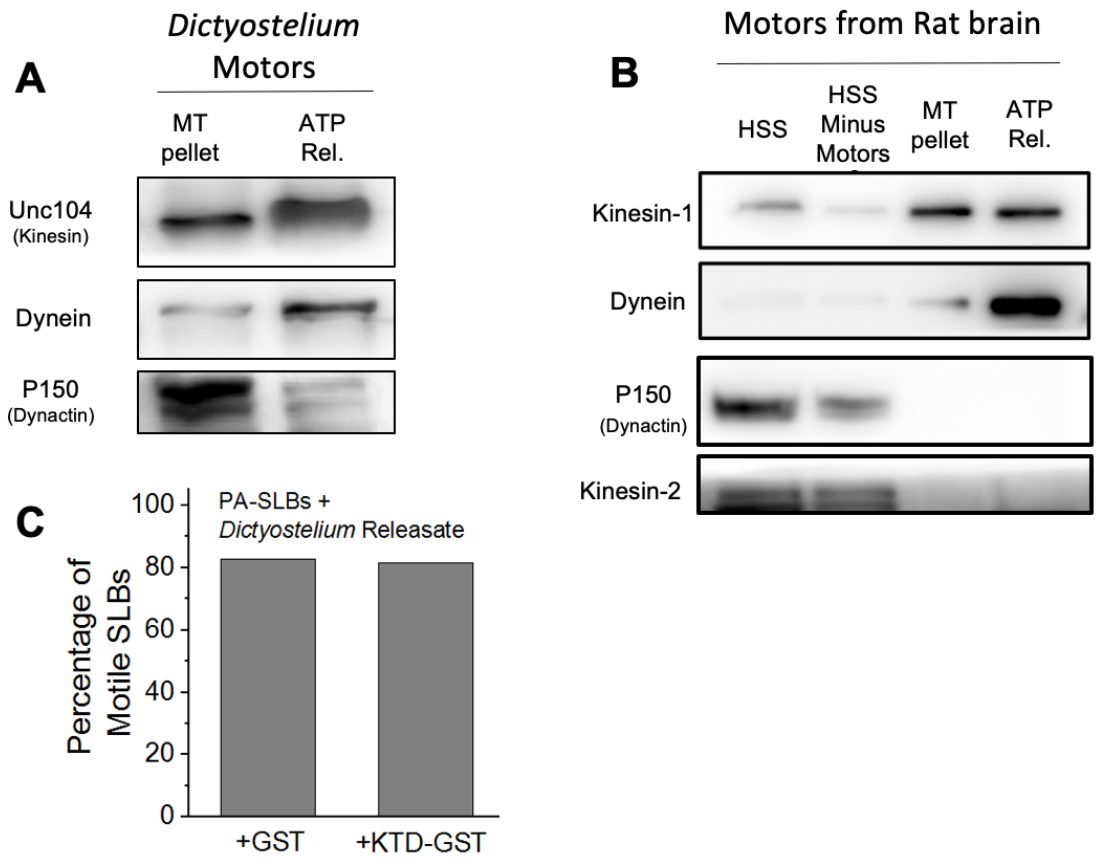


### Supplementary Figure 3

A. Western blots probed for Dynein, Unc104, and Dynactin (P150) in the MT pellet and ATP releasate fractions from *Dictyostelium*. **MT pellet** is the pellet containing MTs after the ATP-release step has been done (see next). **ATP Rel.** is the supernatant after MTs have been pelleted down in the presence of 10mM ATP. ATP releasate is the motor-enriched fraction that is added to SLBs in the motility assay. The MT pellet and ATP releasate lanes contain 75µg total protein per lane.

B. Western blots probed for Kinesin-1, Dynein, Dynactin (P150), and Kinesin-2 in different fractions obtained during the preparation of ATP releasate from rat brain. **HSS** High-speed supernatant (= clarified cytosol) after pelleting the cell membranes. *In-vitro* polymerized MTs are added to this HSS for binding motors and MAPs. **HSS Minus Motors** is the supernatant obtained after MT-bound motors and MAPs are pelleted down from the HSS. Equal volumes of HSS and HSS-Motors were loaded. **MT pellet** is the pellet containing MTs after the ATP-release step has been done (see next). **ATP Rel.** is the supernatant after MTs have been pelleted down in the presence of 10mM ATP. The MT pellet and ATP releasate lanes contain 15µg total protein per lane.

C. Motile fraction of PA-SLBs prepared from *Dictyostelium* releasate in presence of control (GST) and Kinesin Tail Domain (KTD-GST). The kinesin-1 motor is not present in *Dictyostelium*. KTD-GST has no effect on motility of PA-SLBs, demonstrating that KTD-GST is a specific inhibitor of rat-brain kinesin-1 on PA-SLBs, and that it is not a generic inhibitor of Dynein or of other motors. Total of 38 and 52 SLBs prepared with *Dictyostelium* releasate were analyzed respectively for GST and KTD-GST treated groups.

**Phosphatidic Acid Dependent Recruitment of Microtubule Motors to Supported Lipid Bilayers for *In-vitro* Motility Assays,** *Pankaj Kumar et. al.*

#### METHODS

**Antibodies** - Mammalian-DIC (Santacruz Sc13524), Kinesin-1 antibody is described in Ref. (1). Kinesin-2 (Abclonal 11259), P150 (BD Biosciences 612709), Tubulin (Invitrogen 236-10501) The *Dictyostelium*-Dynein antibody was generated against Dynein stalk head, and Unc104 antibody has been described in Ref. (2), *Dictyostelium*-Dynactin antibody was generated against the CC1 domain of dynactin as described (3). Plant-actin antibody was used for detecting *Dictyostelium* actin (Abclonal AC009), GST (Cloud Clone TAX158Ge22). For western blotting, all the antibodies were used at 1:3000 dilution

**Lipids and other chemicals**

All the lipids namely Phosphatidylcholine (PC), phosphatidic acid(PA), Egg sphingomyelin, Cholesterol, Phosphatidyserine (PS), Phosphatidyl-inositol(4,5)bisphosphate ($PIP_2$) and Rhodamine-Phosphatidylethanolamine (Rhod-PE) were obtained from Sigma (Avanti Polar lipds). The GST-KTD peptide was expressed and purified from bacteria (*E. coli*, BL21) as described (1). Ciliobrevin-D was obtained from Sigma. The polycarbonate latex beads (500nm, 900nm, 3μm diameters) were obtained from Polysciences.

**Liposome and SLB preparation**

Phosphatidylcholine (PC; for PC liposomes) or PC and Phosphatidic acid (PA) in the molar ratio 19:1 (for PA liposomes) were dried in a nitrogen stream to form lipid films. A 0.01 molar percentage of Rhodamine-phosphatidylethanolamine (Rh-PE) was added whenever needed to visualize the lipid bilayer membrane on the SLBs. The dried lipid films were stored in a vacuum desiccator for 2 hours. The lipid films were hydrated by vortexing in presence of water to form liposomes at the concentration of 1mg lipids per milliliter of distilled water, and then sonicated in a bath sonicator. These liposomes were used to make the specific types of SLB. Washed beads, 1M NaCl solution, water, and liposomes were incubated in the ratio 5:1:69:20 (ratio of volume in ml) for 30 minutes with intermittent vortexing. The so-formed SLBs were washed twice with water at room temperature for 10 minutes, and then used for assays within the next 10 hours. All spins involving silica beads were carried out at 1000*g* and those for latex beads were carried out at 10,000*g*.

**ATP-Releasate Preparation** - Motor-enriched fractions were prepared as described in reference (4). Briefly, *Dictyostelium* cells or Rat brain tissue were lysed using lysis buffer consisting 30% sucrose (LB-30) along with protease inhibitors. The lysate was given a low-speed (1800g) spin at 4°C for 5 minutes. This was followed by a high-speed spin in an MLS50 rotor (2,25,000g) on LB-30 at 4°C for 30 minutes. A high-speed supernatant (HSS; clarified cytosol) was collected after this spin. 360μg of goat brain tubulin was polymerized using BGT (BRB 80 + 20μM taxol + 10mM GTP). Polymerized microtubules, hexokinase (15U/ml), glucose (4mM), taxol (0.2mM), AMP-PNP (1mM), and $MgCl_2$(4mM) were added to the HSS and incubated on ice for 20 minutes to deplete the ATP. This microtubule-cytosol mixture was centrifuged on LB-30 sucrose gradient at 1,20,000g for 20 min at 4°C. The microtubule pellet was collected and dissolved in the release buffer (LB-15+10mM ATP+10mM $MgCl_2$ with protease inhibitors) for 20-30 mins in ice. This MT-motor mixture was spun at 1,80,000g in a TLA100 rotor for 20 min at 4°C. The supernatant was collected as ATP-releasate, aliquoted and snap snap-frozen in liquid nitrogen.

**Polarity labeling of Microtubules** – Minus-ends of microtubules were labeled as described in (5). Briefly, biotinylated tubulin was polymerized at 37°C for 45 min and sheared by passing through an 18G syringe needle for about 40 times. Small seeds of biotin-tubulin were mixed with unlabeled tubulin in a 1:2 ratio and polymerized at 37°C. Microtubules labeled at the minus end with biotin tubulin were then stuck to poly-lysine coated coverslips and visualized using avidin-coated magnetic beads (150nm diameter).

**Motility assay and Optical Trapping**- Plasma-cleaned coverslips were kept in a Poly-l-lysine solution (1ml Poly-L-Lysine (P8920 Sigma), 36ml ethanol, 4ml methanol) for 30 min and dried in a hot-air oven. Flow chambers are made by sticking these coverslips to a glass slide using double-stick tape (3M; 100 microns thickness). Microtubules were flowed in the chamber and incubated at room temperature for 10 min. The flow chamber was placed a microscope with a custom-built optical trap (6) and microtubules were visualized under Differential Interference Contrast (DIC). Images were acquired at 30 frames/sec. A quadrant photodiode (QPD) was used to record SLB motion. QPD data was digitized at 2 kHz for recording SLB motion. To calibrate the trap, thermal fluctuations of SLBs were recorded at 40 kHz and the corner frequency was determined. Trap stiffness was adjusted between 0.05 pN/nm to 0.15 pN/nm, depending on the experiment.

5μg of ATP releasate and 3μl of SLBs (of desired lipid composition) were incubated on ice for 5 min. For the inhibition experiments 17μM of KTD-GST, or 17μM of GST, or 20μM Ciliobrevin-D, or DMSO was added to this mix and kept on ice for an additional 5 minutes. This mixture, along with 1mM ATP and the ATP regenerating system (20 mM MgCl2, 40 mM creatine phosphate, and 40 U/mL creatine kinase) were flown into the flow chamber. Before adding SLBs, coverslips were blocked with casein-taxol solution. SLBs were trapped in the optical tweezer and placed on a single microtubule using the optical trap for 45 seconds. Free runs were assayed by turning the trap OFF once SLBs bound the microtubule, whereas stalls were obtained by keeping the optical trap ON. If no motion was observed within 45 sec, the SLB was termed as non-motile.

**SLB Pull-down Assay** - 500nm latex beads were used to prepare PC or PA-SLBs. The optical density of a given SLB sample at 500nm wavelength (OD500) was determined. This wavelength was chosen because the Optical density of latex beads exhibits a maximum at 500nm. SLB samples were diluted appropriately to make the OD500 equal, following which equal volumes of these "normalized" samples were used. We have characterized OD500 as a reliable parameter to normalize the number of beads (and therefore of SLBs) across samples. For more details, see (3, 7–9). Such OD-normalized samples (SLBs) have equal amount of lipids and proteins. ATP releasate containing 150μg of proteins was added to the SLBs and incubated at room temperature for 10 minutes with intermittent mixing.

For the SLB pull-down in the presence of inhibitors, 17μM of KTD-GST peptide, or PBS, or 20μM Ciliobrevin-D, or DMSO was added to the SLB-releasate mix and kept at room temperature for 15 minutes. The SLBs were washed twice with water. The supernatant was discarded and the SLB-bound proteins were extracted by boiling the pellet at 95˚C in 0.1% SDS-containing PBS. The beads were removed by pelleting, an equal volume of the supernatant was run in an SDS gel and further processed for immunoblotting.

**Microtubule Pelleting Assay** –ATP-releasate (100μg protein content) prepared from Rat brain was incubated with 17μM KTD-GST, or 17μM GST peptide, or 20μM Ciliobrevin-D or DMSO for 10 minutes on ice. 125μg of *in-vitro* polymerized Taxol-stabilized microtubule was added to this mix and incubated for 20 minutes on ice with intermittent mixing. The ATP releasate has ~5mM ATP. This mix was loaded onto a 25% sucrose (with 20μM Taxol and 1mM GTP) gradient and centrifuged at 90,000g for 30 minutes at 4˚C. The pellet was gently washed using PBS and re-suspended in 45ul 6x SDS dye. Equal volumes of these samples were loaded onto an SDS gel and the separated proteins were investigated using western blotting.

**Lipid-blot Assay** – 5 nano-moles of the specific lipids were spotted onto a nitrocellulose membrane and air-dried. The membrane was blocked using 3% BSA in TBS for 1 hour at RT. The membrane was incubated overnight at 4˚C with 150μg/ml of ATP releasate containing 3% BSA in TBS. The membrane was then washed thrice with 0.1% Tween containing TBS for before probing for bound motors. Primary antibody treatment was followed by secondary antibody. *Dictyostelium*-Dynein antibody (generated against Dynein Stalk Head) and Kinesin-1 antibody were used at 1:4000 for 2 hours at RT in 5% skimmed milk.

**Confocal imaging** – PC-SLBs and PA-SLBs made with liposomes containing 0.01% Rhodamine-Phosphatidylethanolamine (Rhod-PE) in the lipid mixture were imaged on Zeiss 900 upright system using 63X, 1.4-NA objective.

**Data Analysis and Statistics** – Image-J was used to quantify band intensities of western blots and the backgrounds. Origin-2019 was used to prepare the graphs. Custom developed software in LabView (National Instruments) was used to visualize optical trap-generated QPD data. Wherever data was known to be normally distributed, a two-tailed Student's *t-test* was used to test the significance of the results (95% confidence interval).

#### Supplementary Movie Captions

1. **Motion of PC-SLB (*Dictyostelium*)**
   The free-run of a PC-SLB (500nm diameter latex bead) driven by *Dictyostelium* motors on *in-vitro* polymerized microtubules is depicted in the video. The video was acquired at 30fps. The PC-SLB showed frequent pausing between runs.
2. **Motion of PA-SLB (*Dictyostelium*)**
   The free-run of a PA-SLB (500nm diameter latex bead) driven by *Dictyostelium* motors on *in-vitro* polymerized microtubules is depicted in the video. The video was acquired at 30fps. The PA-SLB demonstrate a smoother run.
3. **Motion of PC-SLB (Rat)**
   The free-run of a PC-SLB (500nm diameter latex bead) driven by rat motors on *in-vitro* polymerized microtubules is depicted in the video. The video was acquired at 30fps. The SLB demonstrates a run with intermittent pausing.
4. **Motion of PA-SLB (Rat)**
   The free-run of a PA-SLB (500nm diameter latex bead) driven by rat motors on *in-vitro* polymerized microtubules is depicted in the video. The video was acquired at 30fps.
